# Beyond the foliage: Using non-destructive multimodal 3D imaging and AI to phenotype and diagnose trunk diseases

**DOI:** 10.1101/2022.06.09.495457

**Authors:** Romain Fernandez, Loïc Le Cunff, Samuel Mérigeaud, Jean-Luc Verdeil, Julie Perry, Philippe Larignon, Anne-Sophie Spilmont, Philippe Chatelet, Maïda Cardoso, Christophe Goze-Bac, Cédric Moisy

**Author notes:** These authors contributed equally to this work (Romain Fernandez, Cédric Moisy). Corresponding author: Cédric Moisy.

## Abstract

Quantifying healthy and degraded inner tissues in plants is of great interest in agronomy, for example, to assess plant health and quality and monitor physiological traits or diseases. However, detecting functional and degraded plant tissues *in-vivo* without harming the plant is extremely challenging. New solutions are needed in ligneous and perennial species, for which the sustainability of plantations is crucial. To tackle this challenge, we developed a novel approach based on multimodal 3D imaging and Artificial Intelligence (AI)-based image processing that allowed a noninvasive diagnosis of inner tissues in living plants. The method was successfully applied to the grapevine (*Vitis vinifera* L.) in vineyards where sustainability was threatened by trunk diseases, while the sanitary status of vines cannot be ascertained without injuring the plants. By combining MRI and X-ray CT 3D imaging with an automatic voxel classification, we could discriminate intact, degraded, and white rot tissues with a mean global accuracy of over 91%. Each imaging modality contribution to tissue detection was evaluated, and we identified quantitative structural and physiological markers characterizing wood degradation steps. The combined study of inner tissue distribution *versus* external foliar symptom history demonstrated that white rot and intact tissue contents are key measurements in evaluating vines’ sanitary status. We finally proposed a model for an accurate trunk disease diagnosis in grapevine. This work opens new routes for precision agriculture and *in-situ* monitoring of wood quality and plant health across plant species.

## INTRODUCTION

Wood is a complex biological structure providing physical support and serving the needs of a living plant. Its degradation by stresses or pathogens exposes the plant to huge physiological and structural changes, but the consequences might not be immediately detectable from the outside. Trunk diseases can spread internally and silently, degrading woody tissues, then erratically leading to external symptoms, production losses and, ultimately, the plant’s death. While accurate detection and monitoring of wood degradation require structural and functional characterization of internal tissues, collecting such data on entire plants—nondestructively and *in-vivo—is* extremely challenging. As a result, the study and management of wood diseases are almost impossible *in-situ*. These diseases generate enormous losses on a global scale, including for sectors such as the wine industry.

Grapevine trunk diseases (GTDs) are a major cause of grapevine *(Vitis vinifera* L.) decline worldwide (Claverie et al., 2020). They are mostly undetectable until advanced stages are reached, and the European Union has banned the only effective treatment, i.e., an arsenic-based pesticide. Therefore, vineyard sustainability is jeopardized, with yearly losses of up to several billion dollars (Guerin-Dubrana et al., 2019). Detecting and monitoring GTDs is extremely difficult: fungal pathogens insidiously colonize trunks, leaving different types of irreversibly decayed tissues (Claverie et al., 2020). The predominant GTD, Esca dieback, erratically induces tiger stripe-like foliar symptoms, but their origin remains poorly understood (Bortolami et al., 2019), and their sole observation is not indicative of the vines’ sanitary status (Mugnai et al., 1999; Péros et al., 2008, Lecomte et al., 2012; Maher et al., 2012). Quantifying degraded tissues within living vines could help determine the plant’s condition and predict disease evolution. However, classical techniques (Reis et al., 2019) require sacrificing the plant and often yield limited information. Reaching a reliable diagnosis is thus impossible in living plants.

Monitoring wood degradation using non-destructive imaging techniques has primarily been tested on detached organs, blocks, or planks, and using a single technique: Magnetic Resonance Imaging (MRI) (Pearce et al., 1994; Hiltunen et al., 2020) or X-ray computed tomography (CT) (Van den Bulcke et al., 2009; Hervé et al., 2014; Hamada et al., 2016; Li et al., 2016). In grapevine, CT scanning allowed the visualization of the graft union (Milien et al., 2012), the xylem refilling (Brodersen et al., 2018), tyloses-occluded vessels (Czemmel et al., 2015), and the quantifying of starch in stems (Earles et al., 2018). Synchrotron X-ray CT has been applied to leaves to investigate the origin of foliar symptoms related to trunk diseases, suggesting that symptoms might be elicited from the trunk (Bortolami et al., 2019). X-ray CT and MRI were successfully combined to collect anatomical and functional information to investigate stem flow *in-vivo* (Bouda et al., 2019). These techniques were recently tested for trunk disease detection (Vaz et al., 2020) but were applied separately and on different wood samples, preventing the possibility of combining modalities and thus limiting their effectiveness. They failed to distinguish healthy and defective tissues using MRI.

Imaging-based plant studies are usually performed at a microscopic scale and on a specific detached tissue or organ but rarely on a whole plant. The field of investigation is limited by difficulties in adapting imaging devices to plants and in coupling signals collected from different imaging modalities, preventing the development of ‘digital twin’ models (Laubenbacher et al., 2021) for plants.

Moreover, monitoring wood degradation using multimodal imaging necessitates proper registration and the identification of specific signatures for structural and physiological traits of interest. Such signatures are not well described on wood but are mandatory to deploy automatic quantitative approaches. Experts should perform a preliminary conjoint analysis of multimodal imaging signals together with manual pixel-wise annotation of wood degradation before deploying automatic morphological phenotyping methods based on tissue segmentation and machine learning.

To address these issues, we developed an end-to-end workflow for *in-vivo* phenotyping of internal woody tissues condition, based on multimodal and non-destructive 3D imaging, and assisted by AI-based automatic segmentation. This approach was applied to vine imaging datasets acquired in a clinical imaging facility. 3D images were acquired in five different modalities (X-rays for structure, three MRI parameters for function, and serial sections for expertise) on entire plants, and combined by an automatic 3D registration pipeline (Fernandez and Moisy 2021). Based on serial section annotations, structural and physiological signatures characterizing early and late stages of wood degradation were identified in each imaging modality. Then a machine-learning model was trained to automatically detect tissue condition based on the multimodal imaging data, achieving high performance in recognizing healthy and sick tissues. We could, therefore, perform an accurate quantification of intact, degraded, and white rot compartments within entire vine trunks. We also evaluated the contribution and efficiency of each imaging technique for tissue detection. Finally, we studied the relationships between external foliar symptom expression and the distribution of internal healthy and sick tissues.

This study highlights the potential of our 3D image- and AI-based workflow for non-destructive and *in-vivo* diagnosis of complex plant diseases in grapevine and other plants. It gives access to key indicators to evaluate vines’ inner sanitary status and permits structural and functional modeling of the whole plant. Plant-specific’ digital twins’ (Laubenbacher et al., 2021) could revolutionize agronomy by providing dedicated models for diseased plants and computerized assistance to diagnosis. This work also opens new routes for a broad range of applications on other organs, plants, and diseases, for developing *in-situ* diagnosis solutions, and precision agriculture.

## RESULTS

### Multimodal 3D imaging of healthy and sick tissues

Based on foliar symptom history, *symptomatic-* and *asymptomatic-looking* vines (twelve total) were collected in 2019 from a Champagne vineyard (France) and imaged using four different modalities: X-ray CT and a combination of multiple MRI protocols: T1-, T2-, and PD-weighted (w) (Fig. 1). Following imaging acquisitions, vines were molded, sliced, and each side of the cross-sections photographed (approx. 120 pictures per plant). Experts manually annotated eighty-four random cross-sections and their corresponding images according to visual inspection of tissue appearance. Six classes showing specific colorations were defined (Fig. 2.a): (i) healthy-looking tissues showing no sign of degradation; and unhealthy-looking tissues such as (ii) black punctuations, (iii) reaction zones, (iv) dry tissues, (v) necrosis associated with GTD (incl. Esca and Eutypa dieback), and (vi) white rot (decay). The 3D data from each imaging modality—three MRIs, X-ray CT, and registered photographs—were aligned into 4D-multimodal images (Fernandez and Moisy 2021). It enabled 3D voxel-wise joint exploration of the modality’s information and its comparison with empirical annotations by experts (Fig. 2.b).

**Figure 1:**
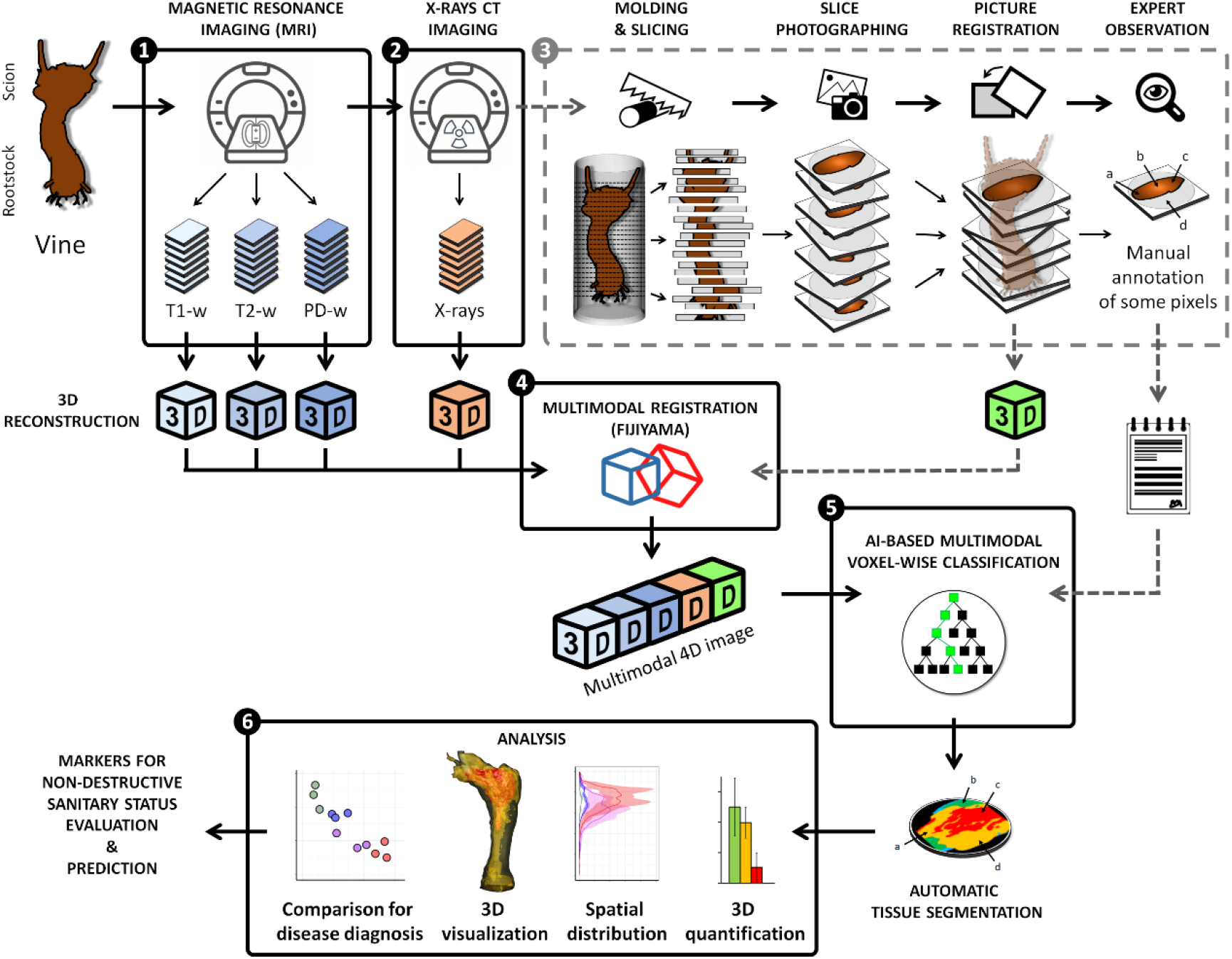
General Workflow: From multimodal vine imaging to data analysis. (1) and (2) Multimodal 3D imaging of a vine using MRI (T1-weighted, T2-w, and PD-w) and X-ray CT. (3) (Optional step) the vine is molded and then sliced every 6 mm. Pictures of cross-sections (both sides) are registered in a 3D photographic volume, and experts manually annotate some cross-sections. (4) Multimodal registration of the MRI, X-ray CT, and photographic data into a coherent 4D image using Fijiyama (Fernandez and Moisy 2021). (5) Machine-learning-based voxel classification. Segmentation of images based on the tissue expert manual annotations: wood *(intact, degraded, white rot), bark*, and *background*. The classifier was trained and evaluated using manual annotations collected on different vines during Step 3. (6) Data analysis and visualization.

**Figure 2:**
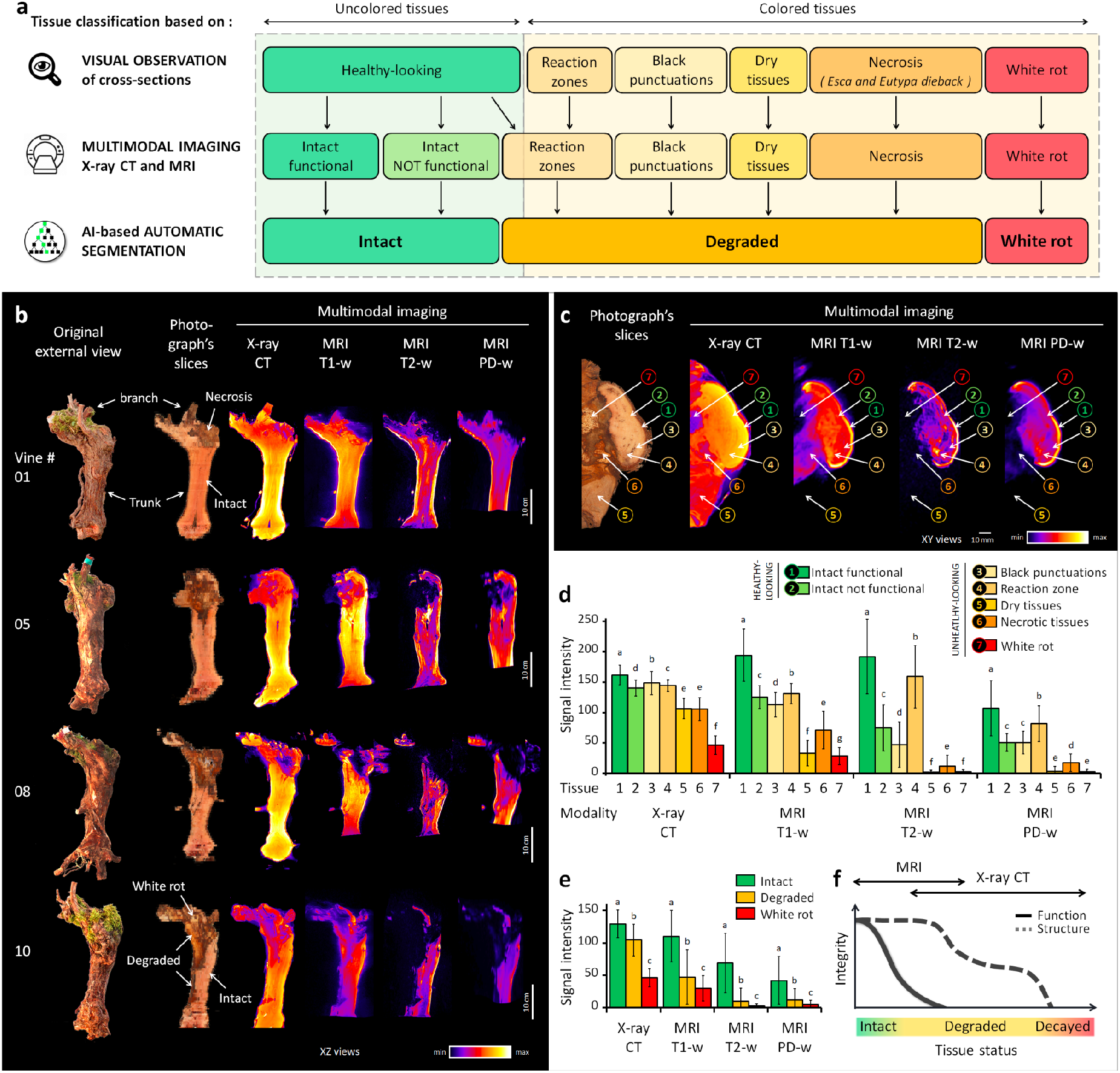
Multimodal imaging and signal analysis. a) Comparison of tissue classification based either on visual observation of trunk cross-sections (6 classes), multimodal imaging data (7 classes), or AI-based segmentation (3 classes). b) Multimodal imaging data collected on vines. XZ views of the photographic, X-ray CT, and MRI volumes, after registration using Fijiyama (Fernandez and Moisy 2021). c) Example of manual tissue annotation and corresponding multimodal signals. d) Multimodal signal values collected by manual annotation of tissues on random trunk cross-sections (19,372 voxels total). e) Multimodal signal values collected automatically on all 4D datasets (46.2 million voxels total) after AI-based voxel classification in three main tissue classes defined as *intact, degraded*, or *white rot*. f) General trends for functional and structural properties during the wood degradation process, and proposed fields of application for MRI and X-ray CT imaging. *Legend: letters on bar plots correspond to Tukey tests for comparing tissue classes in each modality*.

A preliminary study of manually annotated random cross-sections led to the identification of general signal trends distinguishing healthy- and unhealthy-looking tissues (Fig. 2.a, 2.c, and 2.d). In healthy-looking wood, areas of functional tissues were associated with high X-ray absorbance and high MRI values, i.e., high NMR signals in T1-, T2-, and PD-weighted images, while nonfunctional wood showed slightly lower X-ray absorbance (approx. −10%) and lower values in all three MRI modalities (−30 to - 60%).

In unhealthy-looking tissues, signals were highly variable. Dry tissues, resulting from wounds inflicted during seasonal pruning, exhibited medium X-ray absorbance and very low MRI values in all three modalities. Necrotic tissues, corresponding to different types of GTD necrosis, showed medium X-ray absorbance (approx. −30% compared to functional tissues) and medium to low values in T1-w images, while signals in T2-w and PD-w were close to zero (−60 to −85%). Black punctuations, known as clogged vessels, generally colonized by the fungal pathogen *Phaeomoniella chlamydospora*, were characterized by high X-ray absorbance, medium values in T1-w, and variable values in T2-w and PD-w. Finally, *white rot*, the most advanced stage of degradation, exhibited significantly lower mean values in X-ray absorbance (−70% compared to functional tissues; −50% compared to necrotic ones) and in MRI modalities (−70 to −98%).

Interestingly, some regions of healthy-looking, uncolored tissues showed a particularly strong hypersignal in T2-w compared to the surrounding ones (Fig. 2.d). Located in the vicinity of necrotic tissue boundaries and sometimes undetectable by visual inspection of the wood, these regions likely corresponded to the reaction zones described earlier, where host and pathogens strongly interact and show specific MRI signatures (Pearce 2000).

These results highlighted the benefits of multimodal imaging in distinguishing different tissues for their degree of degradation, and in characterizing signatures of the degradation process. The loss of function was correctly highlighted by a significant MRI hyposignal. The necrosis-to-decay transition was marked by a strong degradation of the tissue structure and a loss of density revealed by a reduction in X-ray absorbance. While distinguishing different types of necrosis remained challenging because their signal distributions overlap, degraded tissues exhibited multimodal signatures permitting their detection. Interestingly, specific events such as reaction zones were detected by combining X-ray and T2-w modalities. Overall, MRI appeared to be better suited for assessing functionality and investigating physiological phenomena occurring at the onset of wood degradation when the wood still appeared healthy (Fig. 2.f). In contrast, X-ray CT seemed more suited for discriminating more advanced stages of degradation.

### Automatic segmentation of *intact, degraded*, and *white rot* tissues using non-destructive imaging

To propose a proper *in-vivo* GTD diagnosis method, we aimed to assess vine condition by automatically and nondestructively quantifying the trunks’ healthy and unhealthy inner compartments in 3D. To achieve this complex task, we trained a segmentation model to detect the level of degradation voxel-wise, using imaging data acquired with non-destructive devices. We defined three main classes corresponding to the level of tissue degradation: (1) ‘*intact*’ for functional or nonfunctional but healthy tissues; (2) ‘*degraded*’ for necrotic and other altered tissues; and (3) ‘*white rot*’ for decayed wood (Fig. 2.a).

An algorithm was trained to automatically classify each voxel in one of the three classes based on its T1-w, T2-w, PD-w, and X-ray absorbance values (Fig. 3.a). The classification was performed using the Fast Random Forest algorithm implemented in the Weka library (Witten et al., 2016). The algorithm was first trained on a set of 81,454 manually annotated voxels (Table S1), then cross-validated, and finally applied to whole 4D-images (46.2 million voxels total) (Fig. 1.4). The mean global accuracy of the classifier (91.6% ± 2.0) indicated a high recognition rate, with minor variations among cross-validation folds (Table S2). In our evaluation, F1 scores were 93.6% (± 3.7) for *intact*, 90.0% (± 3.8) for *degraded*, and 91.4% (± 6.8) for *white rot* tissue classes. The global confusion matrix of the validation sets, summed over the 66 folds (895,994 samples), showed that the great majority of incorrect classifications were either due to confusion between *intact* and *degraded* classes (53.4%) or between *degraded* and *white rot* (20.8%) (Table S3). *Intact* and *white rot* classes were seldom confused (<0.001% error).

**Figure 3:**
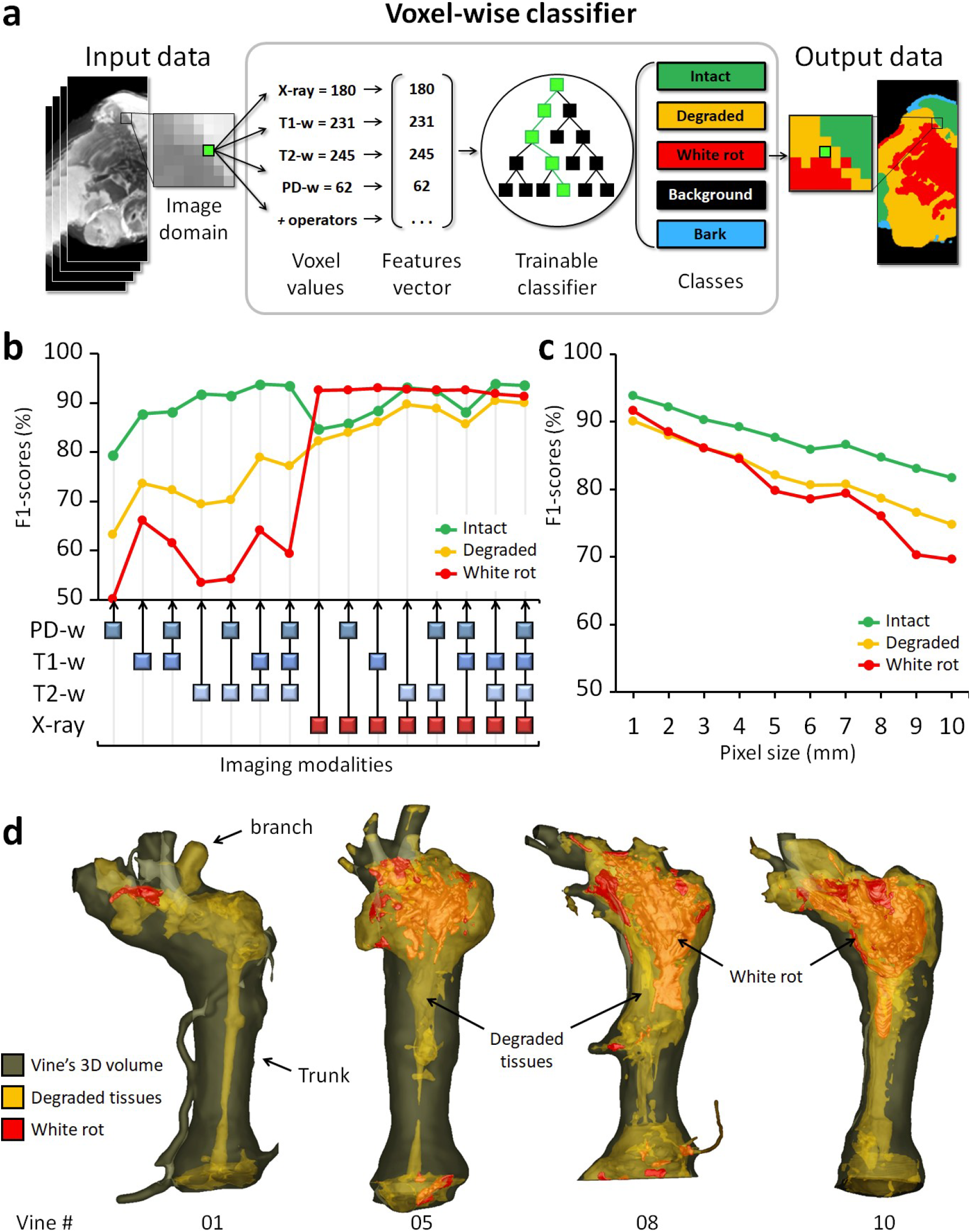
Automatic tissues segmentation. a) AI-based image segmentation using multimodal signals. b) Comparison of all possible imaging modality combinations for their effectiveness (F1-scores) in tissue detection. c) Effectiveness of tissue detection at lower imaging resolutions (using four modalities). d) 3D reconstructions highlighting the extent and localization of the *degraded* and *white rot* compartments in four vines.

The same validation protocol was used to compare the effectiveness of all possible combinations of imaging modalities for tissue detection (Fig. 3.b): the most efficient combination was [T1-w, T2-w, X-ray] for detection of *intact* (F1 = 93.9% ± 3.4) and *degraded* tissues (90.5% ± 3.2); and [T1-w, X-ray] for *white rot* (93.0% ± 5.1). Interestingly, the X-ray modality alone reached almost similar scores for *white rot* detection. In general, slightly better results (± 0.5%) were obtained without considering the PD-w modality, likely due to its lower initial resolution.

The classifier was finally applied to the whole dataset. Statistics were computed to compare the tissue contents in different vines. Considering the entire classified dataset (46.2 million voxels), mean signal values significantly declined between *intact* and *degraded* tissues (−19.3% for X-ray absorbance; and - 57.3%, −86.3% and −71.3% for MRI T1-w, T2-w, and PD-w, respectively); and between *degraded* and *white rot* (−56.0% for X-ray absorbance; and −36.8%, −76.8% and −64.2% for MRI T1-w, T2-w, and PD-w, respectively) (Fig. 2.e and Table S4).

With the increasing deployment of X-ray and NMR devices on phenotyping tasks, in-field imaging has become accessible, but at a heavy cost in terms of image quality and resolution. We challenged our method by training and evaluating the classifier at coarser resolutions, ranging from 0.7 to 10 mm per voxel (Fig. 3.c). Results proved our approach maintained correct performances even at 10 mm (F1 ≥ 80% for *intact*; ≥ 70% for *degraded* and *white rot)*, while a human operator can no longer recognize any anatomical structure or tissue class at this resolution.

These results confirmed the wide range of potential applications and the complementarity of the four imaging modalities. Combining medical imaging techniques and an AI-based classifier, it was possible to segment intact, degraded, and white rot compartments automatically and nondestructively inside the wood. This represents an important breakthrough in their visualization, volume quantification and localization in the entire 3D volume of the vines (Fig. 3.d).

### Deciphering the relationship between inner tissue composition and external symptomatic histories: A step toward a reliable *in-situ* diagnosis

Non-destructive detection of GTDs in vineyards is currently only possible through observing foliar symptoms and vine mortality. Numerous studies are based on these proxies for phenotyping. Foliage is usually screened at specific periods of the year when Esca or Eutypa dieback symptoms occur, and this screening is usually repeated for one or several years. However, new leaves are produced each year, and symptoms may not recur in following years, making any diagnosis hazardous at best, and attempts to correlate external and internal impacts of GTDs unsuccessful. Here, foliar symptoms were recorded yearly for twenty years, since the plot was planted in 1999 (Fig. 4.a). Together with the accurate quantification and localization of degraded and non-degraded compartments in trunks, this allowed more advanced investigations.

**Figure 4:**
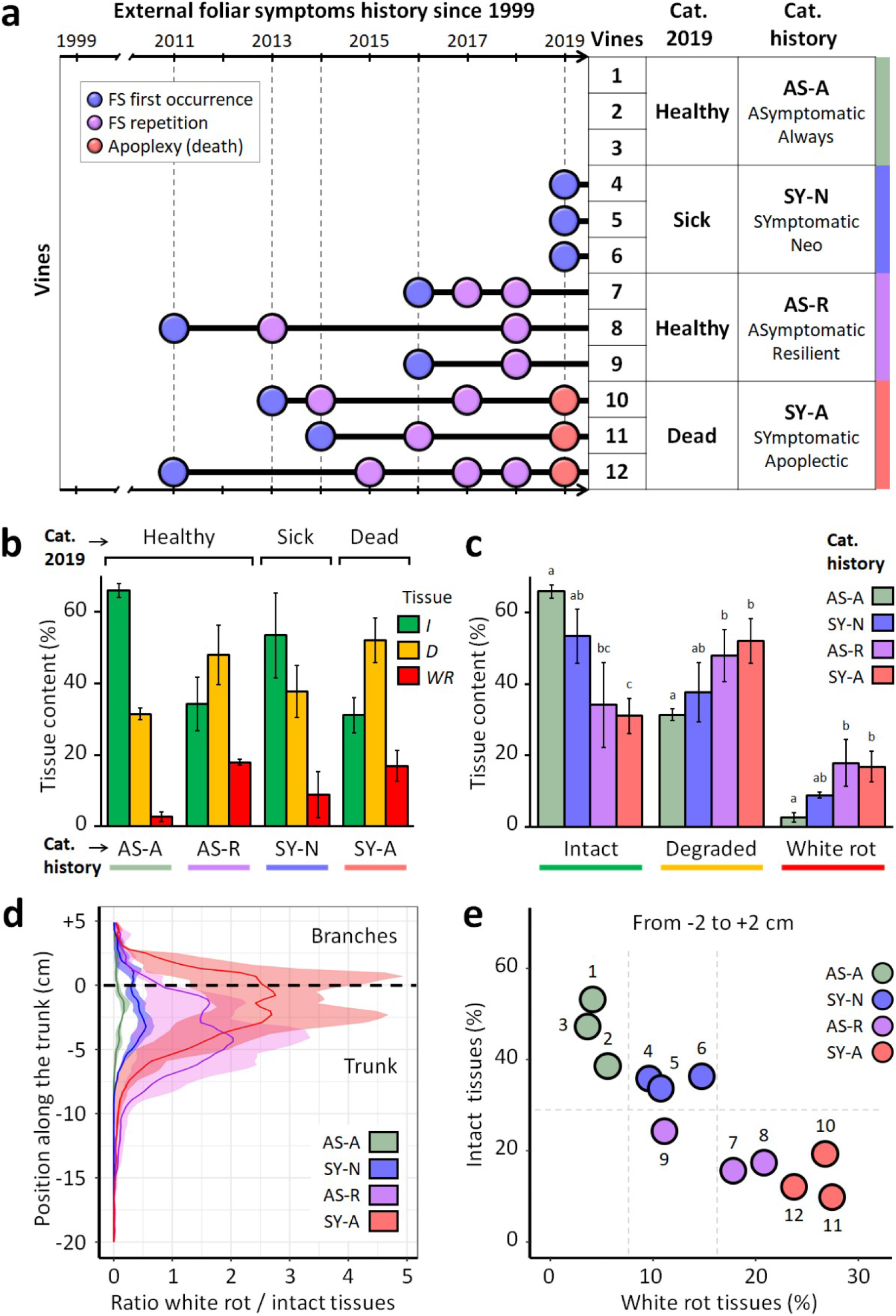
Deciphering the relationship between inner tissue degradation and external foliar symptoms. a) Left: Detailed history of external GTD symptoms expression. Right: classification of vines based on their external sanitary status, either considering year 2019 only or the complete symptom history (right). b) and c) Internal tissue contents of the trunks. Vines are grouped per phenotypic categories, based either on the single 2019 observation or the complete 1999-2019 symptom history. Tissue percentages are calculated from the upper 25 cm of the trunk. d) Comparison of phenotypic categories for the *white rot* and *intact* tissues distribution (mean and interval) along the trunk. Position 0 cm corresponds to the top of the trunk and initiation of branches (i.e., > 0 in branches; < 0 in trunk). e) Comparison of vines for *intact* and *white rot* tissue contents in the region −2 to +2 cm (last 2cm of the trunk and first 2 cm of the branches).

Half of the vines would have been misclassified as *‘healthy’* plants if solely using the foliar symptoms observed in 2019 as markers, despite harboring significantly degraded internal tissues (Fig. 4.b). Indeed, despite the absence of leaf symptoms, these vines contained important volumes of deteriorated wood (up to 623 cm^3^ of *degraded* tissues and 281 cm^3^ of *white rot*) (Table S5). Utilizing the foliar symptom proxy would have led to different -and erroneous-diagnoses each year, confirming its unreliability (Fig. 4.a). Correlations between foliar symptoms observed in 2019 and total internal contents were very weak (R^2^ = −0.25, 0.27, and 0.18 for *intact, degraded*, and *white rot*, respectively). In contrast, internal tissue contents were better supported by categories taking into account the complete vine history (Fig. 4.c). For example, the sum of foliar symptoms detected during the vine’s life was strongly correlated to the composition of internal tissues (R^2^ = −0.87, 0.79, and 0.84 for *intact, degraded*, and *white rot*, respectively) and the correlation between inner contents and the date of the first foliar symptom expression was also high (−0.87 for *intact* and 0.91 for *white rot)* (Table S6).

As illustrated by 3D reconstructions (Fig. 3.d), *degraded* and *white rot* compartments were mostly continuous and located at the top of the vine trunk. This result was consistent with previous reports and the positioning of most pruning injuries that are considered pathways for the penetration of fungal pathogens, causing GTDs (Claverie et al., 2020). However, the distribution and volumes of the three tissue classes helped distinguish different degrees of disease severity (Fig. 4.d). In detail, the tissue content located in the upper last centimeters of the trunk and the insertion point of branches allowed efficient discrimination of the vine condition (Fig. 4.e). On one hand, the proportion of *intact* tissues detected in this region discriminated between vines with mild forms of the disease (*intact* > 30%) and vines at more advanced stages (<30%). On the other hand, the proportion of *white rot* distinguished the healthiest vines *(white rot* < 8%), more affected ones (8 to 15%), and the ones facing critical stages (> 15%). The volume of *degraded* tissues and the positioning of intact and white rot tissues allowed for fine-tuning the diagnosis (Fig. S1).

Noninvasive imaging and 3D modeling offered the possibility to access both internal tissue contents and spatial information without harming the plant (Fig. 5). In vines suffering from advanced stages of trunk diseases, *white rot* tissues were surrounded by *degraded* tissues; thin areas of *intact* functional tissues were limited to the periphery of trunk tops. As illustrated in Fig. 5, abiotic stresses such as a fresh wound can also greatly impact the functionality of surrounding tissues and affect plant survival even more.

**Figure 5:**
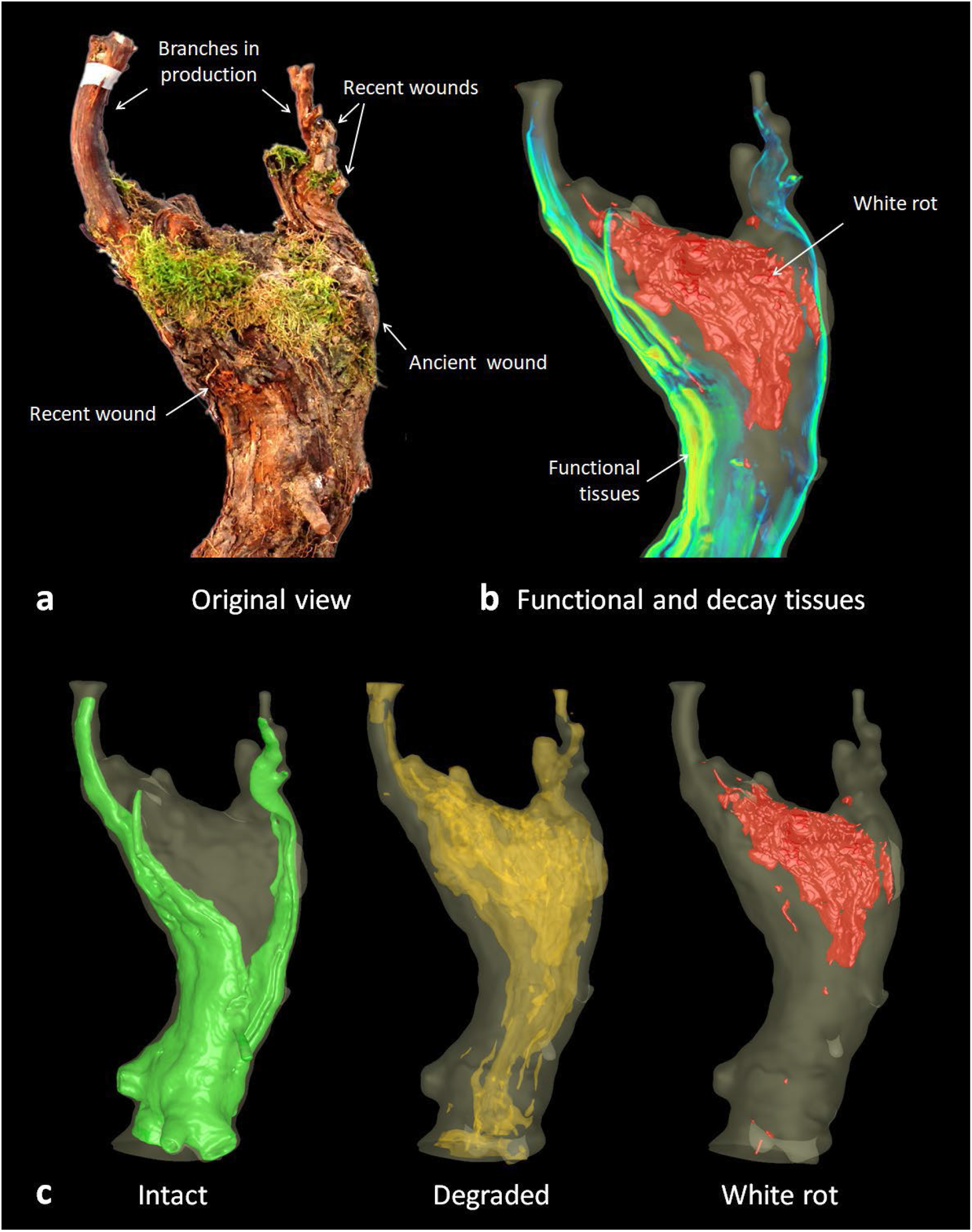
3D visualization of internal tissue contents: example of a specimen at the critical stage of vine decline. a) Original external view of the vine. b) Combining MRI data volume rendering and white rot model. c) 3D representation of the *intact (green), degraded (orange)* and *white rot* (red) compartments inside the trunk.

## DISCUSSION

### A new method for non-destructive detection of wood diseases

Certain plant diseases are mostly undetectable until advanced stages are reached; this is the case for grapevine wood diseases, where their detection is currently only possible through destructive techniques or observing erratically-expressed foliar symptoms. A non-destructive, reliable method for their detection is frequently expected (Gramaje et al., 2018; Mondello et al., 2018; Ouadi et al., 2019; Claverie et al., 2020). To that end, we developed an innovative approach to nondestructively measure healthy and unhealthy tissues in living perennial vines. We combined 1) noninvasive 3D imaging techniques, 2) a registration pipeline for multimodal data, and 3) a machine-learning-based model for voxel classification. We were able to determine the level of tissue degradation voxel-wise, and to accurately segment, visualize, and quantify healthy and unhealthy compartments in the plants.

Among the imaging modalities tested, MRI has already proved relevant in assessing tissue functions in grapevine (Bouda et al., 2019) and in several applications in living plants (Van As and Van Duynhoven 2013). In a recent study, MRI surprisingly failed to distinguish healthy and necrotic tissues in grapevine trunk samples (Vaz et al., 2020). Here, MRI was found particularly well suited for detecting early stages of wood degradation, characterized by a significant loss of signal (57 to 86%) in T1-w and T2-w protocols between *intact* and *degraded* tissue classes. Combining MRI modalities provided information on tissue functionality and water content. T1-w was efficient for anatomical discrimination, and T2-w highlighted phenomena associated with host-pathogens interactions such as reaction zones (Pearce 2000). Interestingly, the T2-w signal dropped by approx. 60% between *functional* and *nonfunctional* tissues but increased by 110% between *nonfunctional* and *reaction zones*. X-ray CT, on the other hand, was particularly efficient in detecting more advanced stages of wood deterioration characterized by a loss in structural integrity, highlighted by a 56% drop in X-ray absorbance between *degraded* and *white rot* tissues.

Multimodal imaging, together with the 4D registration step and machine learning to extract information, proved its efficacy: combining MRI and X-ray CT techniques significantly increased the quality of tissue segmentation. All possible imaging combinations were finally compared for their efficiency, and it is now possible to select the modality/modalities best suited to specific needs. For example, combining T1-w, T2-w, and X-ray was optimal for *intact* and *degraded* tissue detection, but T2-w alone also proved efficient, should using only one imaging modality be possible. For *white rot* detection, combining T1-w and X-ray, or using X-ray alone were the best options.

Given the small size of our training set, we used a strategy based on a random forest classifier, training on the outputs of predefined image processing filters. Though we reached high precision, collecting more data would permit the use of deep learning tools, which could be particularly beneficial in achieving higher levels of predictive accuracy and robustness.

In grapevine, MRI and X-ray CT were recently tested for GTDs detection but applied separately and on different wood samples (Vaz et al., 2020). Here we collected multimodal 3D data on whole trunks of aged plants and developed a pipeline for automatic analysis. Although vines were cut up to gather data to train and evaluate the classifier, this approach is now feasible without harming the plants (Fig. 1). It opens several exciting prospects for diagnosis and applications to other plants and complex diseases.

### GTD indicators based on internal tissue degradation rather than external foliar symptoms

New light was shed on classical monitoring studies when we compared foliar symptom histories with internal degradations. While previous reports showed necrosis volumes could be linked to the probability of esca leaf symptoms occurrence and white rot volumes to apoplectic forms (Péros et al., 2008; Lecomte et al., 2012; Maher et al., 2012; Gramaje et al., 2018), but only weak correlations between tainted or necrotic tissue contents and foliar symptoms were generally observed (Mugnai et al., 1999; Calzarano and Di Marco, 2007). In most studies, plants are considered *‘healthy’* if they do not express any foliar symptoms during the experiment, which generally last just one or two years. Our results confirmed that the appearance of foliar symptoms in a given year is not linked to the volume of internal wood degradation and that a single foliar symptom expression is not a reliable marker of the plant’s actual health status in the GTD context.

Here we considered that the internal tissue composition *(intact, degraded, white rot)* better reflects the severity of the disease affecting the vines and their actual condition. It seemed particularly relevant for *asymptomatic* vines: half of them, harboring large volumes of unhealthy tissues, would have been erroneously categorized as *“healthy”* plants using the foliar symptom proxy. Indeed, *asymptomatic* vines could have reached advanced stages of GTDs, while *symptomatic* ones could be relatively unharmed. In such cases, foliar symptoms-based diagnosis is unreliable; internal tissue content is the only reliable proxy of plant health. Necrotic and decay compartments are intuitively more stable indicators than foliage symptoms: once tissues have suffered irreversible damage, they will clearly remain nonfunctional.

### An internal tissue-based model for accurate GTD diagnosis

A model based on the quality, quantity, and position of internal tissues could be proposed for an accurate diagnosis. Different stages in trunk damage could be distinguished: *‘Low’* damage is characterized by low volumes of altered tissue; ‘*moderate’* by significant *degraded* and decayed contents but still a fair amount of peripheral *intact* tissue; and ‘*critical’* damage, with very limited areas of *intact* tissue (Fig. 6). These stages could be evaluated directly in fields through non-destructive imaging detection, allowing a reliable diagnosis in living specimens.

**Figure 6:**
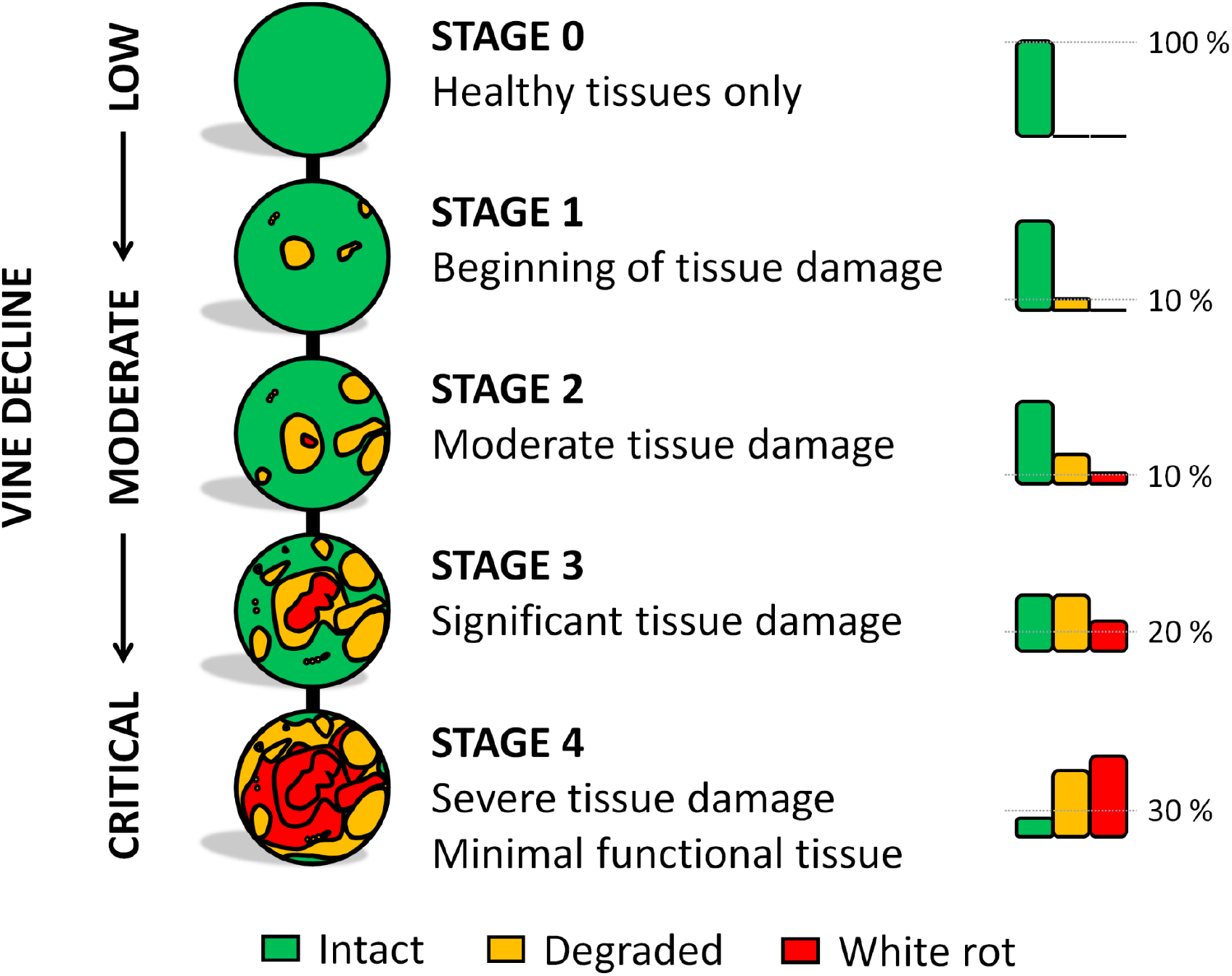
Model for GTD diagnosis based on the degree of trunk internal tissue degradation.

Based on trunk cross-sections, a threshold value of 10% *white rot* in branches has been proposed as a predictor for the chronic form of Esca (Maher et al., 2012; Gramaje et al., 2018). This value could be a proper threshold between the *low* and *moderate* phases defined here, while 20% *white rot* would be the threshold into the *critical* stage. However, *intact* tissues should also be considered: a minimum of 30% *intact* tissues located in the last centimeters of the trunk could be proposed as a threshold for *critical* status.

Here all necrotic tissues were regrouped in a single *degraded* class, but defining more tissue classes— e.g., different types of necrosis—could enable studying each wood disease separately.

Grapevine is a tortuous liana in which both the proportion and configuration of tissues are highly irregular along the trunk and among plants. Thresholds for transitioning from one stage to the other would probably need to be established according to the grapevine variety. Considering their vigor and capacity to produce new functional tissues every year, some varieties might be able to cope with large volumes of unhealthy tissues while maintaining sufficient physiological and hydraulic functions. Other factors, such as the environment, fungal pathogens, and pruning mode, might also influence the plant’s capacity to survive with only very limited *intact* tissues (Claverie et al., 2020), and their impact could be measured using this novel non-destructive approach.

### Predicting the course of diseases and assisting management strategies

The quantity and position of healthy and unhealthy tissues could be useful in predicting the evolution of plants’ sanitary status. It would be tempting to predict that an *asymptomatic* vine might soon develop symptoms according to its *intact* tissue content and assuming its proximity to *neo-symptomatic* vines (Fig. 4.e). Considering *white rot* and *degraded* tissues, we could also estimate that, among the *asymptomatic-resilient* vines, one would be likely to survive a few more years, while others would likely die within a few years. Additional, larger-scale data are required to confirm the effectiveness of these proxies for individual and accurate predictive diagnosis. However, multimodal imaging has already proved relevant for diagnosing the current status of the vines in this study.

White rot removal using a small chainsaw has been proposed to extend the life of seriously affected vines; this technique, *curettage*, currently under evaluation, is a particularly aggressive technique applied ‘blindly’, causing great damage to the plant (Pacetti et al., 2021). A non-destructive imaging-based approach could improve precision surgery by enabling low-damage access to the sick inner compartments by giving access to the exact location and volume of sick tissues to be removed. It will also permit *in-vivo* evaluation of its long-term efficacy.

Finally, non-destructive and *in-vivo* monitoring studies of internal tissue contents could help identify plants that require urgent intervention, i.e., local treatment, curettage, surveillance, or to prioritize replacements in plots, facilitating vineyard management.

## CONCLUSION

By providing direct access to internal tissue degradations in living plants, non-destructive imaging and AI-based image analysis provide new insights into complex diseases affecting woody plants. The results allow for a wide range of new, *in vivo*, and time-lapse studies to become accessible. For example, physiological responses to wounding and pathogen-linked infection could be monitored at the tissue level to search for varietal tolerance. At the individual level, long-term surveillance of healthy, necrotic, and decayed tissues could fine-tune prediction models, permit the evaluation of potential curative solutions, and facilitate the management of agricultural exploitation.

In grapevine, the enigmatic origin of Esca foliar symptoms and the influence of environmental factors on trunk disease development could probably be investigated more efficiently than traditional destructive methods. Based on a limited number of foliar symptom observations, previous studies might also have led to wrong interpretations, and wrong classification of varieties for their tolerance to GTDs, and could be revisited. If no alternative is possible, foliar symptoms should at least be considered with extreme caution after multiple years of surveying.

In medicine, imaging is often dedicated to single individuals, which is rarely the case for plants, which are generally considered at the population level. In viticulture, however, plots are perennial, and each vine represents a long-term financial investment. Individual and non-destructive diagnosis is therefore of great interest for vineyard sustainability, aiding decision-making in targeting a local treatment or replacing specific individuals. Long-term and complex diseases are also generally more difficult to handle.

Conceiving virtual digital twins (Laubenbacher et al., 2021) of living plants would allow for monitoring complex diseases, modeling their evolution, and assessing the impact of novel solutions at different scales. Placing imaging at the bedside of plants offers great hopes and exciting perspectives which could help define next-generation management processes.

## MATERIALS AND METHODS

### Plants

A vineyard was planted in 1999 in Champagne, France, with *Vitis vinifera* L., cultivar Chardonnay rootstock 41B, and uses a traditional Chablis pruning system. Each vine was monitored yearly by CIVC for foliar symptom (FS) expression of GTDs, including Esca, Black dead arm, *Botryosphaeria*, and Eutypa diebacks. Observations were performed at different periods during the vegetative season to optimize the detection of different GTDs, if present.

Based on FS observed in 2019, vines were considered *asymptomatic* (healthy) or *symptomatic* (sick). Based on their whole FS history, vines were then sub-classified as follows:

1. *Asymptomatic-always* if they never expressed any FS.
2. *Asymptomatic-resilient* if they expressed FS in previous years but not in 2019.
3. *Symptomatic-neo* if they expressed FS for the first time in 2019.
4. *Symptomatic-apoplectic* if they died suddenly from typical apoplexy a few days before being collected.

For our study, vines showing different histories—three vines per subclass, 12 total Fig. 3.a)—were manually collected from the vineyard on the 19th of August 2019.

Branches and roots were cut out approximately 15 cm from the trunk, and plants were individually packed in sealed plastic bags to prevent drying.

### Multimodal imaging acquisitions

Multimodal imaging acquisitions were performed on each vine, from rootstocks to the beginning of branches, by Magnetic Resonance Imaging (MRI) and X-ray Computed Tomography (CT). MRI acquisitions were performed with Tridilogy SARL (http://www.tridilogy.com) and the help of radiologists from CRP/Groupe Vidi at the Clinique du Parc (Castelnau-le-Lez, France), using a Siemens Magnetom Aera 1,5 Tesla and a human head antenna. Three acquisition sequences, T1-weighted(-w), T2-w, and PD-w were performed on each specimen, respectively:

1. 3D T1 Space TSE Sagittal (Thickness 0.6 mm, DFOV 56.5 x 35 cm, 320 images, NEx 1, EC 1, FA 120, TR 500, TE 4.1, AQM 256/256).
2. 3D T2 Space Sagittal (Thickness 0.9mm, DFOV 57.4 x 35.5 cm, 160 images, NEx 2, EC 1, FA 160, TR 1100, TE 129, AQM 384/273).
3. Axial Proton Density Fat Sat TSE Dixon (Ep 5mm, Sp 6.5, DFOV 57.2 x 38 cm, 40 images, NEx 1, EC 1, FA 160, TR 3370, TE 21, AQM 314/448).

X-ray CT acquisitions were performed at the Montpellier RIO Imaging platform (Montpellier, France, http://www.mri.cnrs.fr/en/) on an EasyTom 150kV microtomograph (RX Solution). 3D volumes were reconstructed using XAct software (RX solution), resulting in approximately 2500 images per specimen at 177 μm/voxel resolution. Geometry, spot, and ring artifacts were corrected using the default correction settings when necessary.

### Plant slicing and photographic acquisition

After MRI and X-ray CT acquisitions, plants were individually placed in rigid PVC tubes, molded in a fast-setting polyurethane foam filler, and cut into 6 mm-thick cross-sections using a bandsaw (Fig. 1.3). Cutting thickness was approx. 1 mm. Marks were placed on tubes to ensure regular slicing, and three rigid plastic sticks with different diameters were molded together with vines to serve as landmarks for their realignment. Both faces of each cross-section were then photographed using a photography studio, artificial light, a tripod, a digital camera (Canon 500d), and a fixed-length lens (EF 50mm f/1.4 USM) to limit aberration and distortion. Approximately 120 pictures per plant were collected and registered into a coherent 3D photographic volume based on landmarks.

### Data preprocessing: 4D multimodal registration

For each vine, 3D data from all modalities (MRI T1-w, T2-w and PD-w, X-ray CT, and 3D photographic volumes) were registered using Fijiyama (Fernandez and Moisy, 2021) and combined into a single 4D-multimodal image (voxel size = 0.68 mm x 0.68 mm x 0.60 mm) (Fig. 1.4). To compensate for possible magnetic field biases, which are generally present at the edge of the fields during MRI acquisitions, checkpoints were added manually, facilitating the estimation of non-linear compensations during 4D registration.

The registration accuracy was validated using manually placed landmarks (167 couples) distributed in the different modalities. Compared to MRI and X-ray CT modalities, the photographic volume presented a reduced number of images and light geometric distortions due to slicing irregularities. However, the alignment between photographs and other modalities resulted in an estimated average registration mismatch of 1.42 ± 0.98 mm (mean ± standard deviation). The alignment between photographs and other modalities was accurate enough to allow experts to manually annotate tissues directly on the 4D-multimodal images (see below).

### Preliminary investigation of multimodal signals

An initial signal study was conducted on eighty-four cross-sections randomly sampled from three vines. Tissues were firstly classified into 6 different classes based on their visual appearance (Fig. 2.a): (i) *healthy-looking tissues* showing no sign of degradation; (ii) *black punctuations* corresponding to clogged vessels; (iii) *reaction zones* described earlier (Pearce, 2000); (iv) *dry tissues* resulting from pruning injuries; (v) *degraded tissues* including several types of necrotic tissues; and (vi) *white rot*. Once considering X-ray CT and MRI images, experts could distinguish *intact functional* and *intact nonfunctional* tissues among the *healthy-looking* class, resulting in seven tissue classes in total (Fig. 2.a and 2.c). Moreover, some *healthy-looking* tissues showed specific MRI hypo- or hyper-signals and were re-classified as *reaction zones* (Fig. 2.c). For these particular classes, an alteration of the wood aspect was not always visible by direct observation of the cross-sections.

Finally, multiple regions of interest (ROIs, 19,372 voxels total) were delineated by hand on the multimodal images and assigned to one of the seven tissue classes. For each selected voxel, values were gathered simultaneously from the four modalities (X-ray CT, T1-w, T2-w, and PD-w; 77,488 values total) using the registered multimodal images. The data were processed using R (v3.5.3) and the R-studio interface (v1.2.5001). Results are summarized in Fig. 2.d. The significance of differences observed between the seven tissue classes was tested within each modality using Tukey tests and a 95% family-wise confidence level.

### Automatic tissues segmentation of the whole 3D datasets

For each plant, thirteen cross-sections were sampled and manually annotated to label the corresponding voxels. Five classes were defined: background, bark, intact tissues, degraded tissues, and white rot (Fig. 2.a). The annotation was performed using the Trainable segmentation plugin for Fiji (Arganda-Carreras et al., 2017), which was extended to process multi-channel 3D images (see code availability). As a result, a set of 81,454 annotated voxels distributed among the twelve 3D volumes was produced (Table S1).

We trained an algorithm to classify each voxel *P_i_* = (*x,y,z*) of the images (20 million voxels per specimen), attributing a class *C_i_* among the five previously described (Fig 3.a). The classification was performed using the Fast Random Forest algorithm implemented in the Trainable Segmentation plugin, given its performance when working with ‘small’ training datasets (< 100.000 samples).

For each voxel, a feature vector *X_i_* was built and then used by the classifier to predict the class Ĉ_ii_ of the voxel *P_i_*. Information on the voxel’s local environment was gathered in the feature vector by applying various image processing operators, e.g., local mean value, variance, edge, and others, to the initial images and for each imaging modality. These operators were parameterized using a scale factor, taking values from 1 to 64 voxels.

### Evaluation of classifier performances

The classifier performances were evaluated using a k-fold cross-validation strategy. In each fold, the annotated voxels were split into a training set and a validation set. The train set, regrouping annotations from 10 plants, was used to train the classifier. The validation set, containing annotations from the two remaining plants, was used to assess the performances of the trained classifier. A global confusion matrix was computed from all 66 folds (Table S3). This matrix evaluated global and class-specific accuracies and F1-scores—considering both the test precision p and the recall r (Chinchor 1992)—for each class and all possible combinations of imaging modalities (Fig. 3.b and Table S2). F1-scores are generally considered a better indicator of performance because they highlight under- and over-estimations of a specific class more precisely.

### Tissue quantification and 3D volumes reconstruction

For further analysis, we only considered voxels corresponding to areas of interest, i.e., tissue classes *intact, degraded*, and *white rot*. The number and localization of these voxels were collected for tissue quantification and visualization. 3D views presented in Fig. 3.d and Fig. 5 were produced using isosurface extraction and volume rendering routines from VTK libraries (Schroeder and Martin 2005).

### Relative positioning of tissue classes along the vines

To compare the position of *intact, degraded*, and *white rot* tissues in different vines, we estimated the geodesic distance separating each voxel from a common reference area point located at the center of the trunk, twenty centimeters below the trunk head. Using geodesic distances, we considered a region ranging from the last 20 cm of the trunk (defined as “position −20”), passing through the top of the trunk (‘0’), to the first 5 cm of branches (‘+5’) (Fig. S2). This computation allowed the identification of voxel populations located within the same distance range while considering the tortuous shape of the trunks.

### Simulation of performances at lower resolutions

Test images were built by image sub-sampling to simulate an average portable imaging device’s resolution, resulting in voxel sizes ranging from 0.7 (original resolution) up to 10 mm. The corresponding annotated samples were converted accordingly, retaining the most represented label for each voxel volume. The classifier was then trained and tested on these low-resolution sample sets.

## ACKNOWLEDGEMENTS

This work was supported by the French Ministry of Agriculture and Food, France AgriMer, the Comité National des Interprofessions des Vins à appellation d’origine et à indication géographique (CNIV), and the Institut Français de la Vigne et du Vin (IFV) within VITIMAGE and VITIMAGE-2024 projects (program Plan National Dépérissement du Vignoble); and by Agropolis fondation-APLIM Etendard project (contract 1504-005).

Imaging acquisitions were performed at Tridilogy SARL / Groupe CRP, and the MRI platform member of the national infrastructure France-BioImaging supported by the French National Research Agency «Investments for the Future» (ANR-10-INBS-04), and of the Labex CEMEB (ANR-10-LABX-0004) and NUMEV (ANR-10-LABX-0020).

The authors warmly thank Stéphane Bottalico and Renaud Lebrun (CNRS) for technical assistance, and Lindsay Hartley-Backhouse (Scriptoria Solutions) for copy editing.

## AUTHOR CONTRIBUTION

**Romain Fernandez:** Methodology, Software, Investigation, Formal analysis, Visualization, Writing. **Loïc Le Cunff:** Conceptualization, Methodology, Investigation. **Samuel Mérigeaud:** Conceptualization, Methodology, Investigation, Data Curation. **Jean-Luc Verdeil:** Conceptualization, Methodology, Investigation, Writing. **Julie Perry:** Resources, Investigation. **Philippe Larignon:** Investigation, Data Curation, Writing. **Anne-Sophie Spilmont:** Conceptualization, Writing. **Philippe Chatelet:** Investigation, Writing. **Maïda Cardoso:** Conceptualization, Writing. **Christophe Goze-Bac:** Conceptualization, Writing. **Cédric Moisy:** Conceptualization, Methodology, Investigation, Formal analysis, Visualization, Funding acquisition, Supervision, Writing.

## DATA AND CODE AVAILABILITY

The datasets generated and analyzed during the current study are available from the corresponding author upon reasonable request.

The extension of the Trainable Segmentation plugin is open-source, and available as a fork of Trainable Segmentation on GitHub: https://github.com/Rocsg/Trainable_Segmentation/tree/Hyperweka.

## SUPPORTING INFORMATION

The following supporting information is available for this article:

**Table S1.** Number of annotated samples available for classifier training and evaluation. Distribution among tissue classes and vines.

| Vine # | Classes |  |  |  |  | Total samples | % |
| --- | --- | --- | --- | --- | --- | --- | --- |
|  | Background | Intact tissues | Degraded tissues | White rot | Bark |  |  |
| **01** | 683 | 3,749 | 2,699 | 155 | 560 | **7,846** | 9.6 |
| **02** | 781 | 3,079 | 2,490 | 337 | 599 | **7,286** | 8.9 |
| **03** | 252 | 2,231 | 2,106 | 49 | 528 | **5,166** | 6.3 |
| **04** | 538 | 2,290 | 3,155 | 1,450 | 550 | **7,983** | 9.8 |
| **05** | 135 | 2,013 | 2,690 | 2,681 | 192 | **7,711** | 9.5 |
| **06** | 272 | 2,168 | 2,821 | 1,250 | 301 | **6,812** | 8.4 |
| **07** | 317 | 2,743 | 2,343 | 1,169 | 451 | **7,023** | 8.6 |
| **08** | 461 | 2,179 | 2,247 | 665 | 319 | **5,871** | 7.2 |
| **09** | 288 | 2,868 | 1,500 | 479 | 559 | **5,694** | 7.0 |
| **10** | 403 | 2,154 | 2,578 | 2,167 | 180 | **7,482** | 9.2 |
| **11** | 192 | 2,004 | 2,755 | 894 | 358 | **6,203** | 7.6 |
| **12** | 274 | 1,123 | 2,733 | 2,009 | 238 | **6,377** | 7.8 |
| **Total** | **4,596** | **28,601** | **3,0117** | **13,305** | **4,835** | **81,454** | **100** |
| **%** | 5.6 | 35.1 | 37.0 | 16.3 | 5.9 | 100 |  |

**Table S2.**
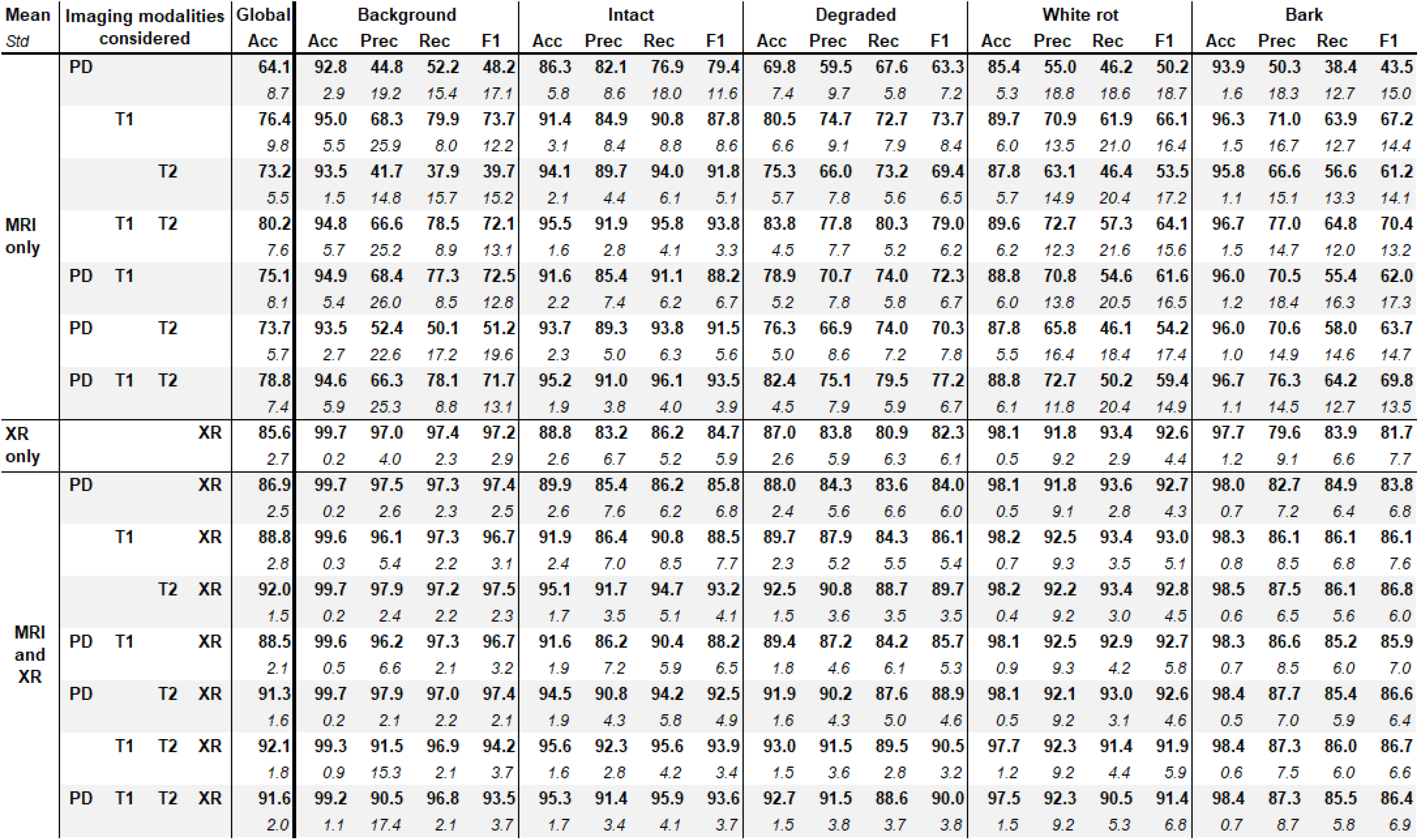
Evaluation of classifier performances. Global and class accuracies (Acc), precision (Prec), recall (Rec) and F1-scores (F1) percentages. Mean (bold) and standard deviation *(italic)* were collected by training on ten vines and evaluating on the last two. Different combinations of imaging modalities were tested: MRI PD-w, T1-w, and T2-w; and X-ray CT (XR).

**Table S3.**
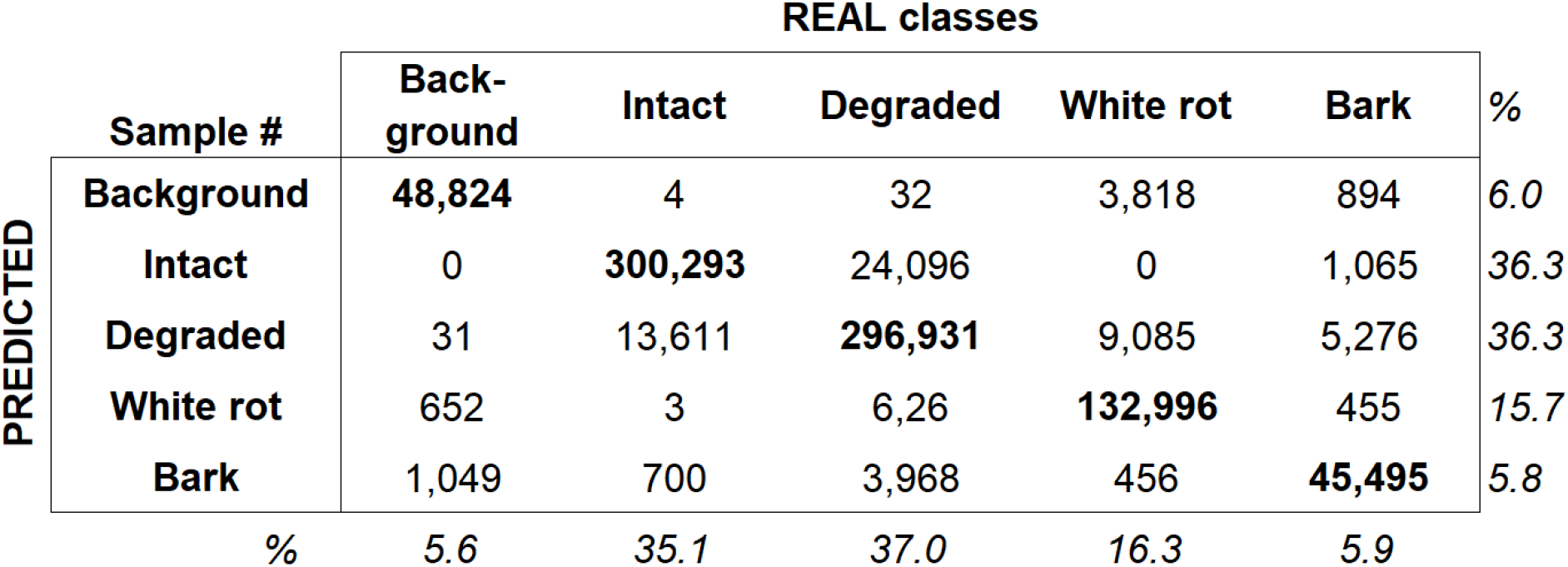
Evaluation of the classifier performance: sum of confusion matrices. Considering the 66 folds of the cross-validation.

**Table S4.** Multimodal signal values corresponding to the three main tissue classes. Means (in bold) and standard deviations (italic) (values in 8-bits) collected on the whole dataset (after automatic classification, 46.2 million voxels total).

| Imaging modalities | Tissue classes |  |  |
| --- | --- | --- | --- |
|  | Intact | Degraded | White rot |
| **X-ray** | **129,4** | **104,5** | **46,0** |
|  | *21,3* | *24,6* | *14,1* |
| **T1-w** | **110,2** | **47,1** | **29,7** |
|  | *40,0* | *42,6* | *19,7* |
| **T2-w** | **68,9** | **9,4** | **2,2** |
|  | *46,1* | *20,4* | *3,0* |
| **PD-w** | **41,8** | **12,0** | **4,3** |
|  | *37,2* | *17,4* | *6,8* |

**Table S5.** Tissue contents per vine. Contents measured for each individual vine from the automatically segmented 3D datasets. Data were collected in the region ranging from the upper last 20 cm of the trunks to the first 5 cm of the branches. Results are expressed as volume (cm3) and percentages.

| Vine # | Phenotype | Tissue content cm<sup>3</sup> (%) |  |  |
| --- | --- | --- | --- | --- |
|  |  | Intact | Degraded | White rot |
| 01 | AS-A ASymptomatic Always | 687 (2,0) | 302 (29,9) | 21 (68,0) |
| 02 | AS-A ASymptomatic Always | 571 (4,2) | 278 (31,3) | 38 (64,3) |
| 03 | AS-A ASymptomatic Always | 663 (1,7) | 336 (33,0) | 18 (65,1) |
| 04 | SY-N Symptomatic Neo | 722 (9,5) | 395 (31,9) | 118 (58,4) |
| 05 | SY-N Symptomatic Neo | 426 (7,9) | 450 (47,2) | 76 (44,7) |
| 06 | SY-N Symptomatic Neo | 559 (9,1) | 334 (33,9) | 90 (56,8) |
| 07 | AS-R ASymptomatic Resilient | 278 (19,2) | 623 (55,8) | 215 (24,9) |
| 08 | AS-R ASymptomatic Resilient | 357 (23,6) | 551 (46,3) | 281 (30,0) |
| 09 | AS-R ASymptomatic Resilient | 421 (10,8) | 368 (41,5) | 96 (47,5) |
| 10 | SY-A SYmptomatic Apoplectic | 333 (20,8) | 436 (44,9) | 202 (34,2) |
| 11 | SY-A SYmptomatic Apoplectic | 425 (12,2) | 685 (54,1) | 155 (33,5) |
| 12 | SY-A SYmptomatic Apoplectic | 252 (17,5) | 565 (57,0) | 174 (25,4) |

**Table S6.**
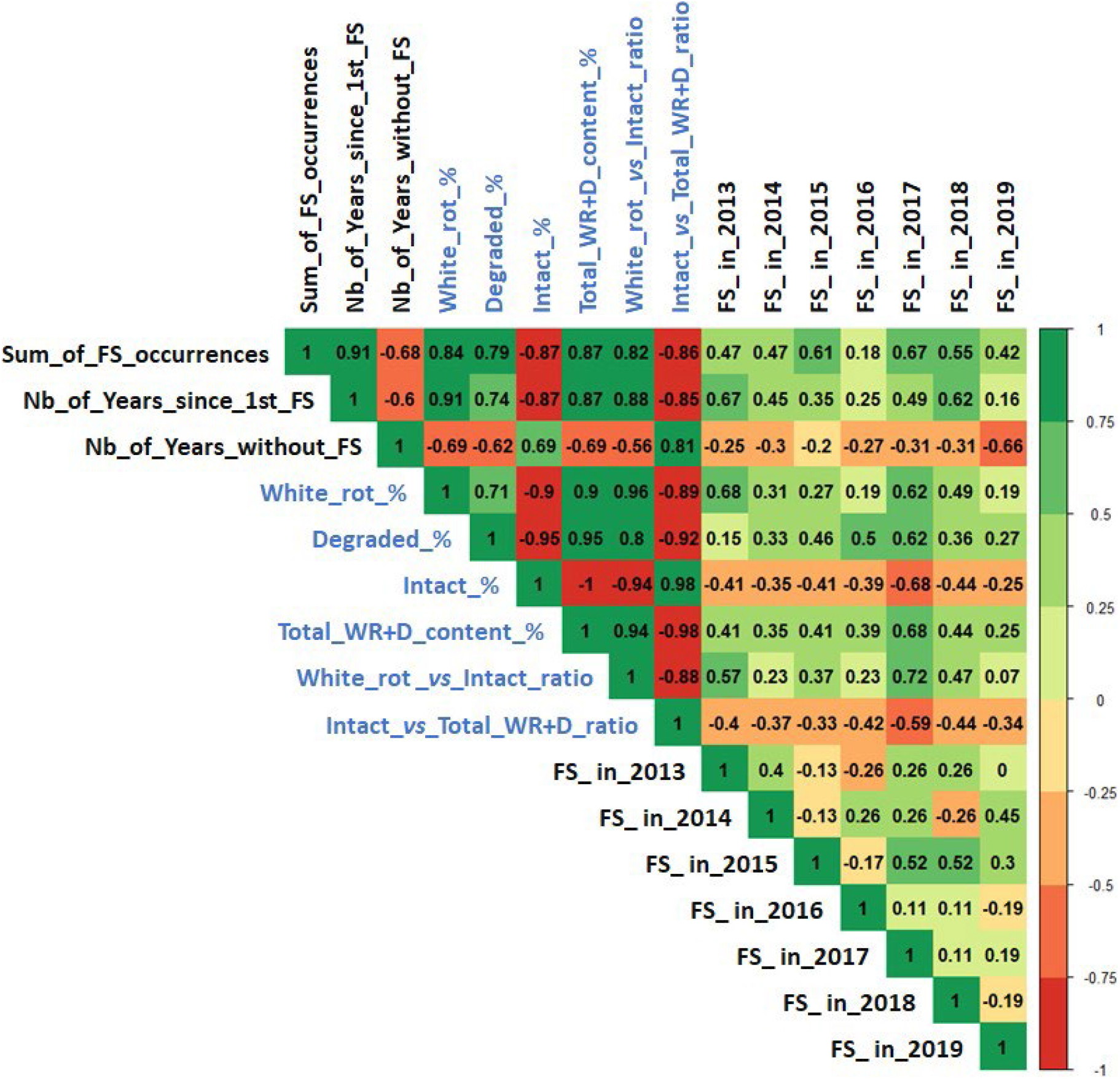
Correlogram. Chart of correlation statistics between “internal” (blue text) and “external” (black text) proxies for GTD status diagnosis. FS = foliar symptom; Nb = number; D = degraded tissues; WR = white rot.

**Fig. S1.**
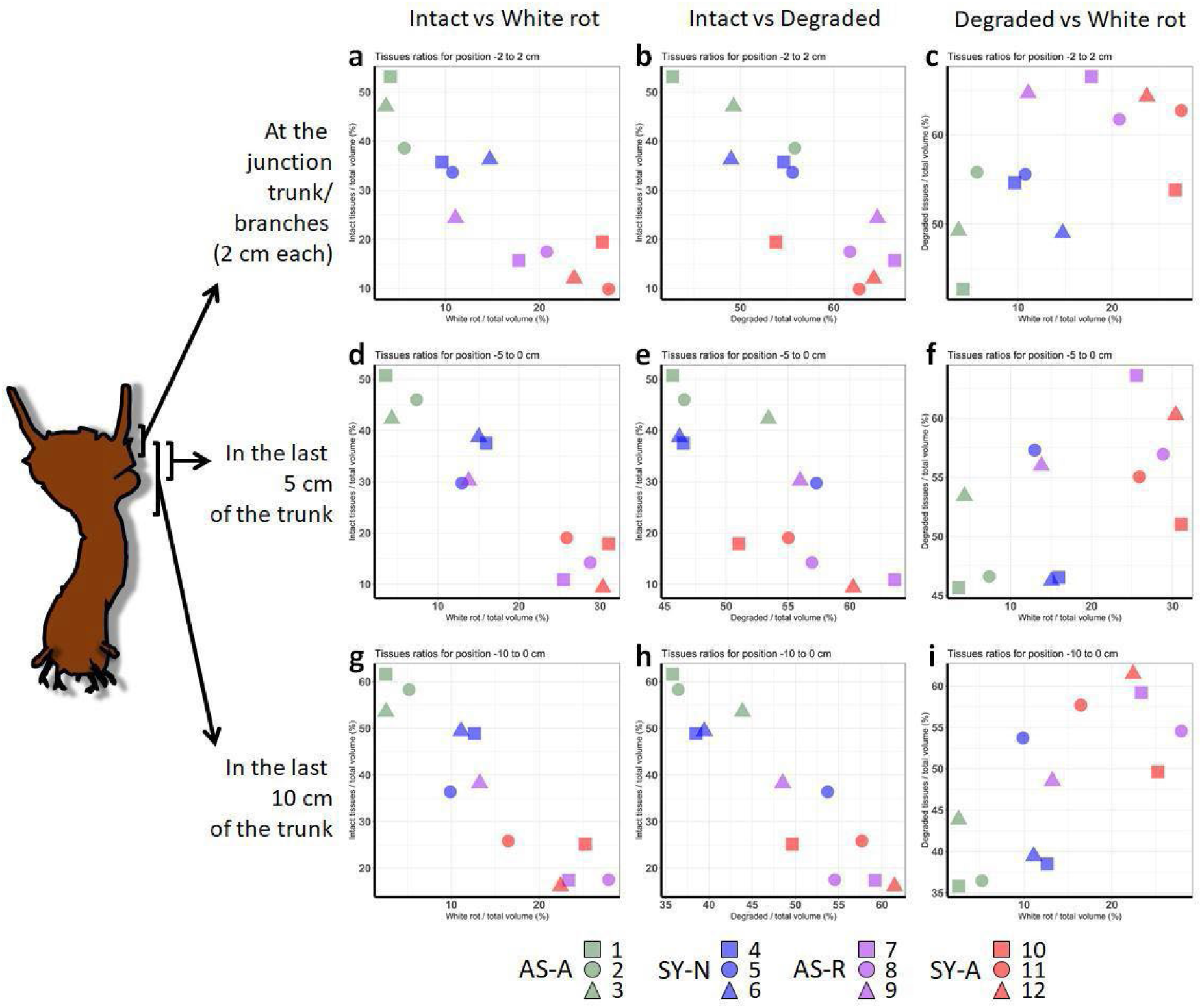
Detailed comparison of vines for intact, degraded, and white rot contents considering different positions along the vine trunk. AS-A = asymptomatic-always; SY-N = symptomatic-neo; AS-R = asymptomatic-resilient; SY-A = symptomatic-apoplectic.

**Fig. S2.**
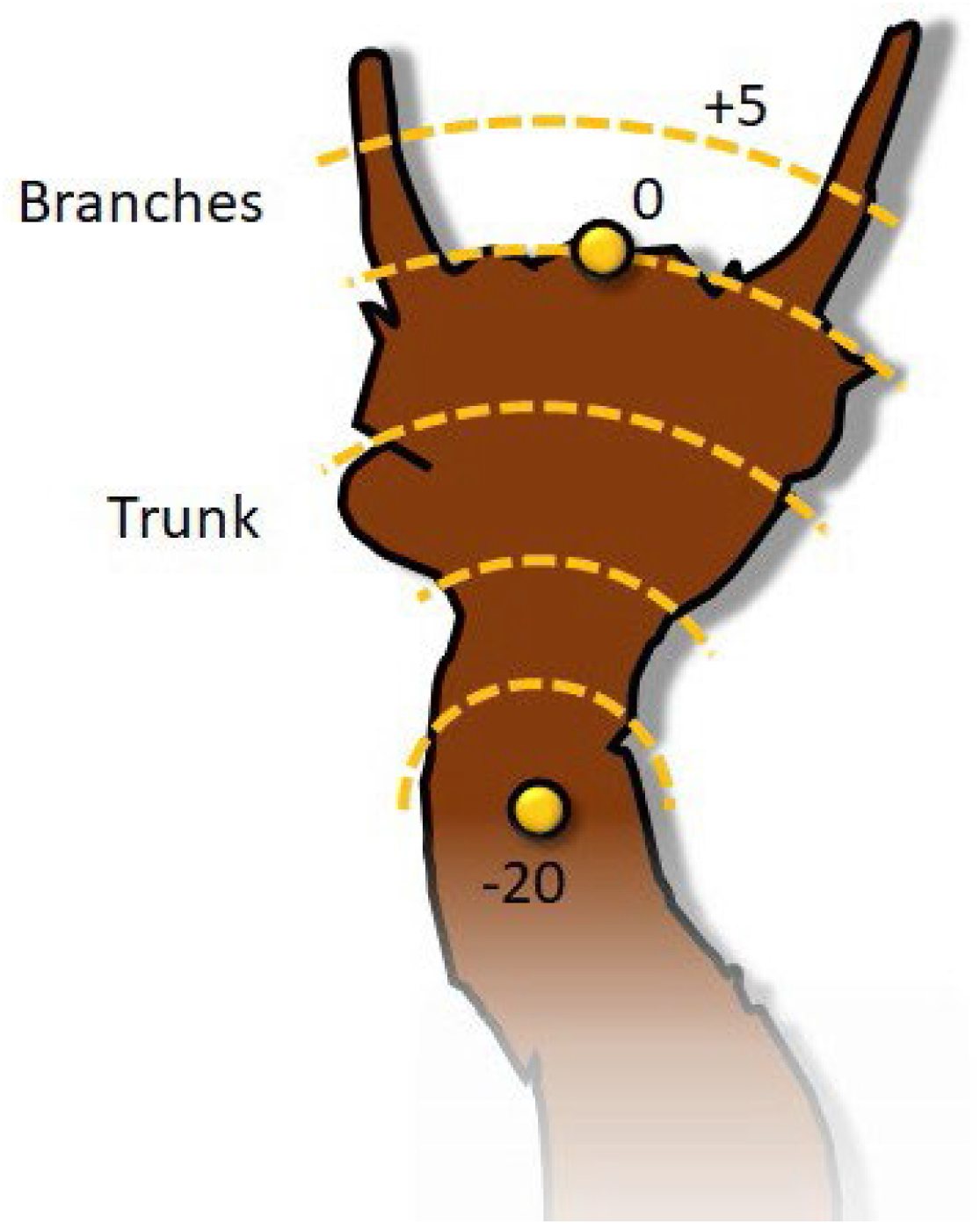
Vine trunk geodesic distance estimation. Geodesic distances were estimated from the center of the trunk and using the top of the trunk as a reference (point “0”).

## Notes

### Competing Interest Statement

The authors have declared no competing interest.

### Summary of Updates

update manuscript

